# Concurrent assessment of gait kinematics using marker-based and markerless motion capture

**DOI:** 10.1101/2020.12.10.420075

**Authors:** Robert Kanko, Elise K. Laende, Elysia M. Davis, W. Scott Selbie, Kevin J. Deluzio

## Abstract

Kinematic analysis is a useful and widespread tool used in research and clinical biomechanics for the estimation of human pose and the quantification of human movement. Common marker-based optical motion capture systems are expensive, time intensive, and require highly trained operators to obtain kinematic data. Markerless motion capture systems offer an alternative method for the measurement of kinematic data with several practical benefits. This work compared the kinematics of human gait measured using a deep learning algorithm-based markerless motion capture system to those of a common marker-based motion capture system. Thirty healthy adult participants walked on a treadmill while data were simultaneously recorded using eight video cameras (markerless) and seven infrared optical motion capture cameras (marker-based). Video data were processed using markerless motion capture software, marker-based data were processed using marker-based capture software, and both sets of data were compared. The average root mean square distance (RMSD) between corresponding joints was less than 2.5 cm for all joints except the hip, which was 3.6 cm. Lower limb segment angles indicated pose estimates from both systems were very similar, with RMSD of less than 5.5° for all segment angles except those that represent rotations about the long axis of the segment. Lower limb joint angles captured similar patterns for flexion/extension at all joints, ab/adduction at the knee and hip, and toe-in/toe-out at the ankle. These findings demonstrate markerless motion capture can measure similar 3D kinematics to those from marker-based systems.

## 1. Introduction

Kinematic analysis is the measurement of the motion of rigid bodies and is an important tool in clinical and research biomechanics. While rigid body kinematics are ubiquitous in biomechanics research, the experimental and computational methods used to estimate three-dimensional (3D) segment position and orientation (pose) vary, and so do the resulting kinematics. Markerless motion capture has the potential to alleviate some of the technical and practical issues of marker-based motion analysis without sacrificing data quality by replacing physical palpation of bony landmarks with probabilistic estimation of segment pose by highly trained neural networks (Mathis et al., 2018).

In traditional marker-based motion analysis, expertise is required to identify anatomical landmarks through physical palpation. This skill necessitates knowledge of anatomy and the translation of this information to a specific individual, a process which is honed over years of experimental work. Lack of skill in this domain leads to the misrepresentation of anatomical landmarks and unrepresentative anatomical coordinate systems and subsequent joint centres, which propagate to errors in reported joint kinematics and kinetics (Della Croce et al., 1999; Holden and Stanhope, 1998; Stagni et al., 2000). This has been noted as one of the greatest sources of error in motion analysis, being greater than instrument error and similar to skin movement artefact (Della Croce et al., 2005). Systematic bias in landmarks for a given examiner may also exist which make the consolidation of different datasets and multi-center collaborations challenging (Gorton et al., 2009; Johnson et al., 2018). In the case of markerless motion analysis, landmark detection is performed by a highly trained deep neural network that applies rules consistently across individuals, thereby dissociating the tracking of human motion from the operator.

Pose estimation in both marker-based and markerless systems is achieved by using at least three noncollinear landmarks to define the pose of body segments, which is represented by a local coordinate system (LCS) relative to a global coordinate system (GCS). In a marker-based system, joint center positions are estimated using regression approaches based on anthropometric measurements from medical imaging or cadaveric studies (Harrington et al., 2007), or functional approaches which approximate the center of rotation between two segments (Ehrig et al., 2006; Hicks and Richards, 2005; Schwartz and Rozumalski, 2005). A markerless system estimates the probability of joint center locations directly from an annotated training set of hundreds of thousands of images. Decision rules for joint center positions are estimated in each frame of data by a highly trained neural network based only on the training the network received and the video recorded of the movement.

Reducing the reliance on the expert by removing physical markers increases the ease of collecting data. Not only does this have the potential to improve the reliability of data (Kanko et al., 2021a), but also to expand the use of quantitative human movement data to areas where attaching physical markers to a participant is impractical, or an impediment to the research itself. Such instances include when a participant is not comfortable wearing minimal and skin-tight clothing, during movements where sweat reduces marker adhesion, and where observation bias overwhelms the research such as behavioral studies (Hutchinson et al., 2019). Markerless motion capture has the potential to measure the kinematics of human movement with fewer limitations than marker-based motion capture, including the ability to collect data outside of a laboratory environment.

The aim of this study was to compare full-body 3D kinematics of gait measured by a deep neural network-based markerless motion capture system to those measured by a marker-based motion capture system.

## 2. Methods

### 2.1 Reuse of Data

The dataset used in this study, which is made up of simultaneous marker-based motion capture, synchronized video, and ground reaction forces from an instrumented treadmill for thirty healthy adult participants during treadmill gait at self-selected speeds, has been previously reported on in a comparison of spatiotemporal gait parameter measurements using marker-based and markerless motion capture systems (Kanko et al., 2021b). This study reused the marker trajectory data, synchronized video data, and ground reaction force data. However, this study differs from the previous comparison in the following ways: the video data were processed using a newer version of the markerless motion capture software, producing separate markerless motion capture data; this study compares kinematic waveforms rather than spatiotemporal parameters, which are often of interest to differing core audiences; the data analysis described here, including gait event detection, was performed independently from that previously described, with the two being treated as two fully separate studies.

### 2.2 Markerless Motion Capture

*Theia3D* (Theia Markerless Inc., Kingston, ON, Canada) is a deep learning algorithm-based approach to markerless motion capture that uses synchronized video data to perform 3D human pose estimation, and which has been previously described in detail elsewhere (Kanko et al., 2021b, 2021a). Briefly, the system utilizes deep convolutional neural networks which were trained on over 500,000 manually annotated digital images of humans in the wild to estimate the position of 51 salient features on humans in new images provided to the system. Using this approach, the system estimates the 2D positions of these features for all humans that appear within synchronized and calibrated video data provided to the system, from which 3D position estimates can be obtained. An articulated multi-body model is scaled to fit the subject-specific landmarks positions in 3D space, and a multi-body optimization approach (inverse kinematic (IK)) is used to estimate the 3D pose of the subject throughout the recorded physical task. By default, the lower body kinematic chain has six degrees-of-freedom (DOF) at the pelvis and three DOF at the hip, knee, and ankle. This markerless system has previously been shown to measure comparable spatiotemporal gait parameters to marker-based motion capture and a pressure-sensitive gait mat, and measure gait kinematics over multiple sessions with greater reliability than marker-based motion capture (Kanko et al., 2021b, 2021a).

### 2.3 Participants

A convenience sample of thirty healthy, recreationally active individuals (15 male/15 female, mean (SD) age: 23.0 (3.5) years, height: 1.76 (0.09) m, mass: 69.2 (11.4) kg) were recruited to participate in this study at the Human Mobility Research Laboratory (Kingston, ON). Participants gave written informed consent, and this study was approved by the institutional ethics committee. Exclusion criteria included having any neuromuscular or musculoskeletal impairments that could prevent their performance of walking. Participants were provided with minimal, skin-tight clothing, and wore their personal athletic shoes. Retroreflective markers and tracking clusters were affixed to relevant anatomical landmarks and body segments; a complete description of the marker set has been previously reported (Kanko et al., 2021b)

### 2.4 Experimental Setup and Procedure

The data collection setup and procedure were previously used to compare spatiotemporal gait parameter measurements between marker-based and markerless motion capture (Kanko et al., 2021b). Specifically, a marker-based camera system (seven Qualisys 3+ (Qualisys AB, Gothenburg, Sweden), 1.3 Megapixel resolution motion capture cameras) and a markerless camera system (eight Qualisys Miqus, 3 Megapixel resolution video cameras) were positioned around an instrumented treadmill (Tandem Force-Sensing Treadmill, AMTI Inc., MA). Both camera systems were connected to a single instance of Qualisys Track Manager for synchronization and to allow them to be calibrated simultaneously, resulting in a single shared global reference frame. Both systems recorded at 85 Hz.

A static calibration trial for the marker-based motion capture data was collected with the subject standing on the treadmill; no static trial is required for the markerless system. Starting at an initial speed of 1.2 m/s, participants determined a comfortable self-selected walking speed by providing feedback to researchers. Participants acclimatized to the treadmill for two minutes before ten consecutive trials of four seconds were collected simultaneously using both camera systems.

### 2.5 Data Analysis

Markerless motion capture video data were processed using *Theia3D* (v2021.1.0.1450), from which 4×4 pose matrices of each body segment were exported for analysis in Visual3D (C-Motion, USA) alongside the tracked marker trajectories from the marker-based motion capture. Two skeletal models were created in Visual3D; one that tracked the markerless pose matrices, which Visual3D created automatically when data from *Theia3D* was loaded, and a second that tracked the marker trajectories, which was manually defined. The marker-based model was defined to have identical joint constraints as those of the *Theia3D* IK model. Force-based gait events were used to obtain time-normalized gait cycles, all of which were included in the analysis except those with tracking issues in the data from either system, which were excluded for both systems; this ensured the measurements being compared between systems were for identical gait cycles. Kinematic measures were calculated using the Visual3D models, time normalized over the gait cycle, and analyzed further in MATLAB (MathWorks, Natick, MA). Joint position estimates, lower limb segment angles, and lower limb joint angles were directly compared for matching gait cycles between systems, which were summarized as subject-average differences and average root-mean-square (RMS) differences across all subjects. Global segment angles represent the components of the XYZ Cardan angle taken between the segment LCS and the laboratory GCS (defined as Z-Up and Y-Anterior); thus, the x-component represents rotation about the global x-axis, the z-component represents rotation about the axially-aligned axis of the segment LCS, and the y-component represents rotation about the floating axis produced by the cross product of the latter with the former. Joint angles were calculated using a Cardan sequence equivalent to the Joint Coordinate System (Grood and Suntay, 1983).

## 3. Results

Data from all thirty subjects were included in the analysis, using an average of 37 gait cycles per subject (range 11 to 51). The majority of identified tracking issues that led to the exclusion of a gait cycle were found in the marker-based data. Example skeletal models from the two systems are overlaid in Figure 1A for visual comparison, and the markerless skeletal model is shown overlaid on the original video data in Figure 1B. Visually, there is substantial overlap between the two models, providing face validity that the pose estimates were similar between the two systems.

**Figure 1:**
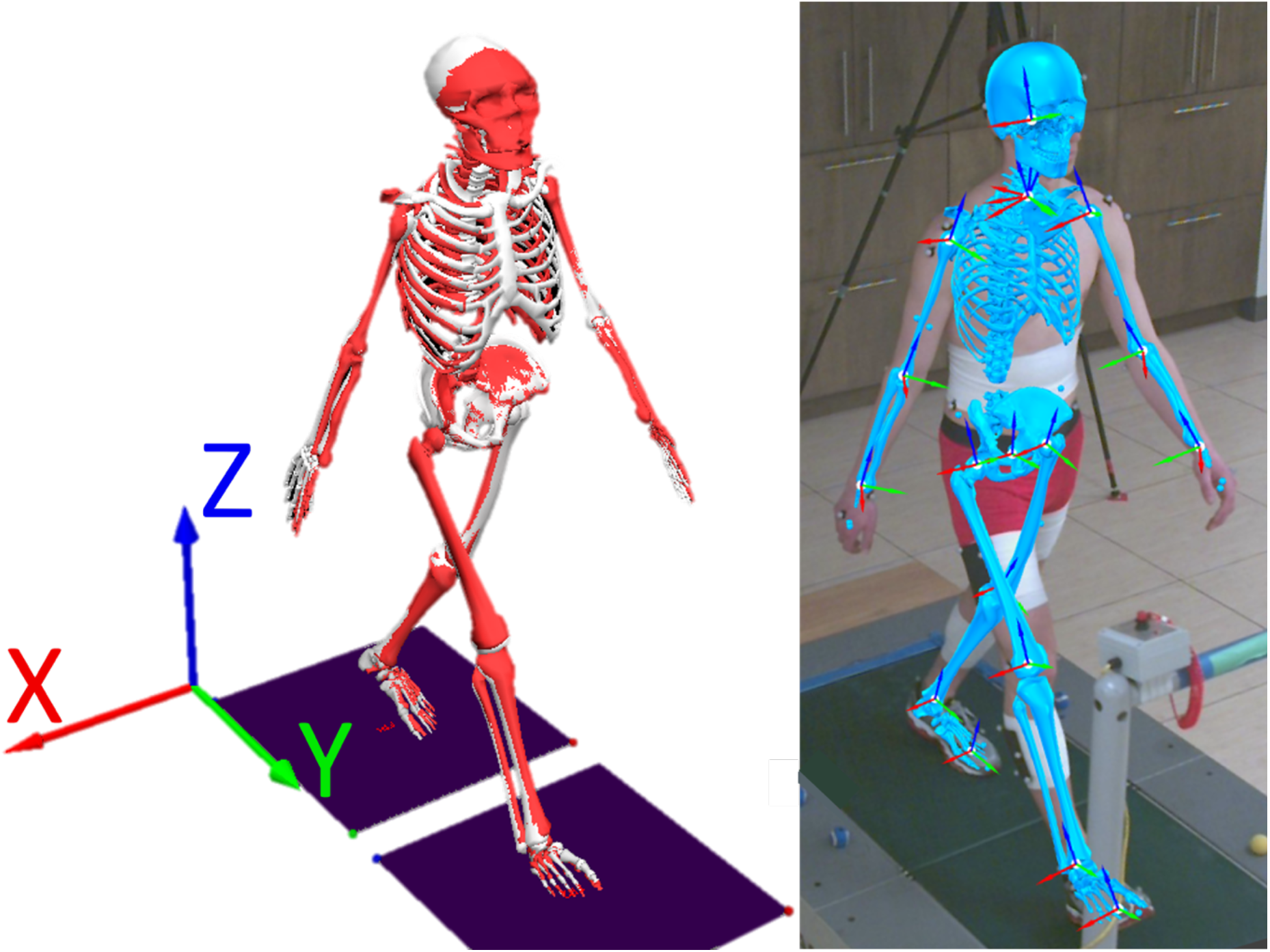
Examples of A) the concurrent skeletal models with the global coordinate system, and B) the markerless skeletal model overlaid on original video data, with segment local coordinate systems displayed. The markerless skeletal model is shown coloured red in the image with the concurrent models, and coloured blue in the overlaid image. The global coordinate system is fixed in the treadmill, with the x-axis pointing laterally, the y-axis pointing in the direction of progression, and the z-axis pointing vertically. This subject had average joint position 3D differences at the ankle, knee, hip, shoulder, elbow, and wrist of 2.6 cm, 1.5 cm, 3.3 cm, 2.2 cm, 1.9 cm, and 1.1 cm, respectively. The joint angle RMS differences between the markerless and marker-based models for this subject were 10.1° in ankle flex/ext, 2.3° in knee flex/ext, 6.5° in hip flex/ext, 6.2° in ankle inv/ever, 3.6° in knee ab/ad, 2.1° in hip ab/ad, 9.8° in foot toe-in/-out, 13.9° in knee int/ext rotation, and 3.1° in hip int/ext rotation angles.

### 3.1 Joint Position Estimates

The average 3D Euclidean distance between corresponding lower limb joints throughout the gait cycle across all subjects was 2.4 cm, 2.2 cm, and 3.6 cm, for the ankle, knee, and hip, respectively. The lower limb joint position differences demonstrated some dependence on gait cycle phase, indicating that there are systematic differences in the estimates from both systems that are dependent on subject pose (Figure 2). The average 3D Euclidean distance between corresponding upper limb joints throughout the gait cycle across all subjects was 2.1 cm, 2.4 cm, and 1.1 cm at the shoulder, elbow, and wrist, respectively. The upper limb joint position estimates were less dependent on gait cycle phase than the lower limb joints, with less variable differences throughout the gait cycle (Figure 3).

**Figure 2:**
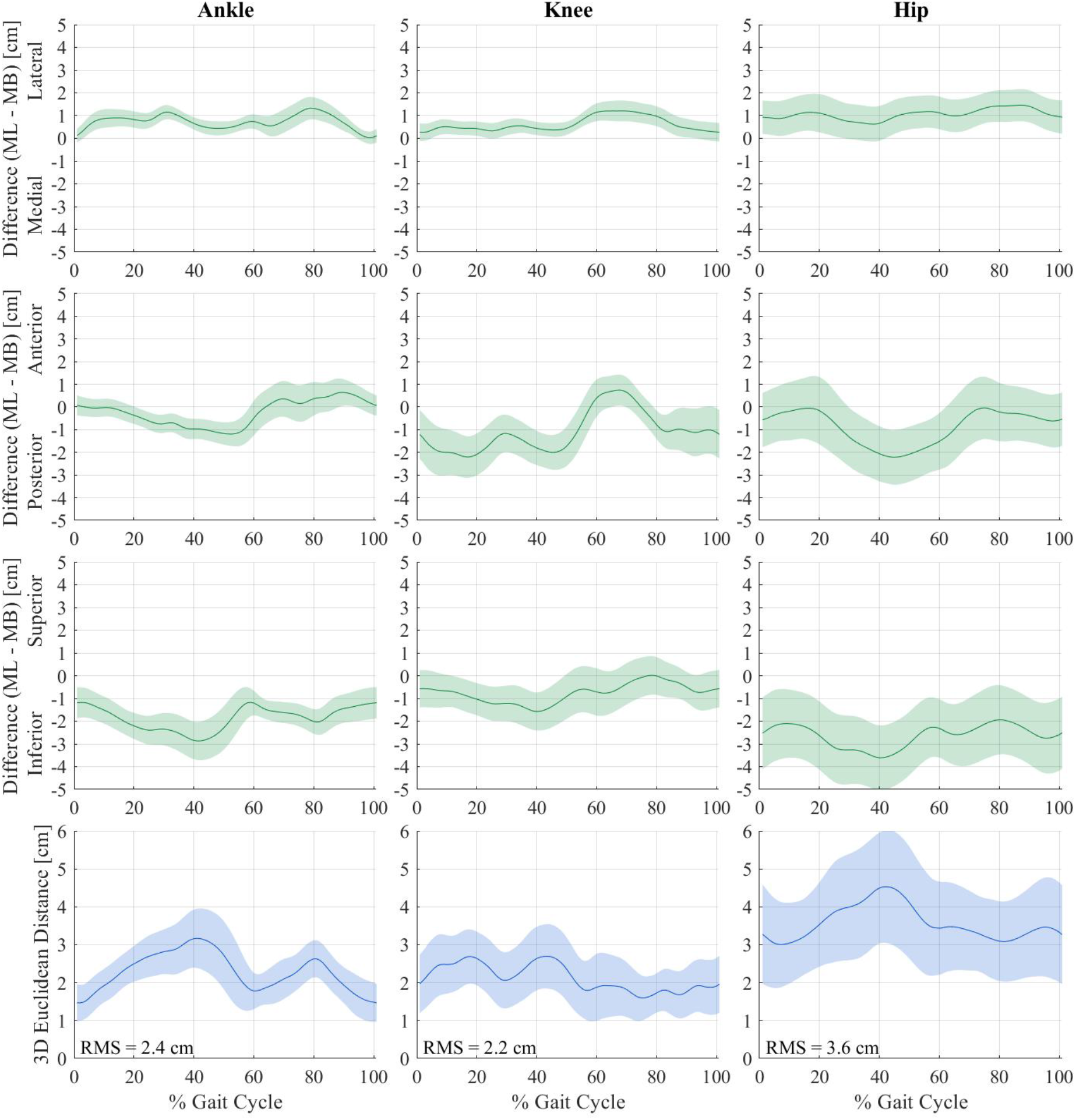
Mean +/− SD lower limb joint position differences between both motion capture systems across the gait cycle for thirty subjects. Differences were calculated as [markerless (ML) position] - [marker-based (MB) position] and are expressed as components along each of the global coordinate system axes (row 1: medial/lateral direction; row 2: anterior/posterior direction; row 3: superior/inferior direction) and as 3D Euclidean distances (row 4). Axis labels (e.g. Lateral) indicate the position of the markerless joint relative to the marker-based joint. Root-mean-square (RMS) values are inset in the 3D Euclidean distance plots.

**Figure 3:**
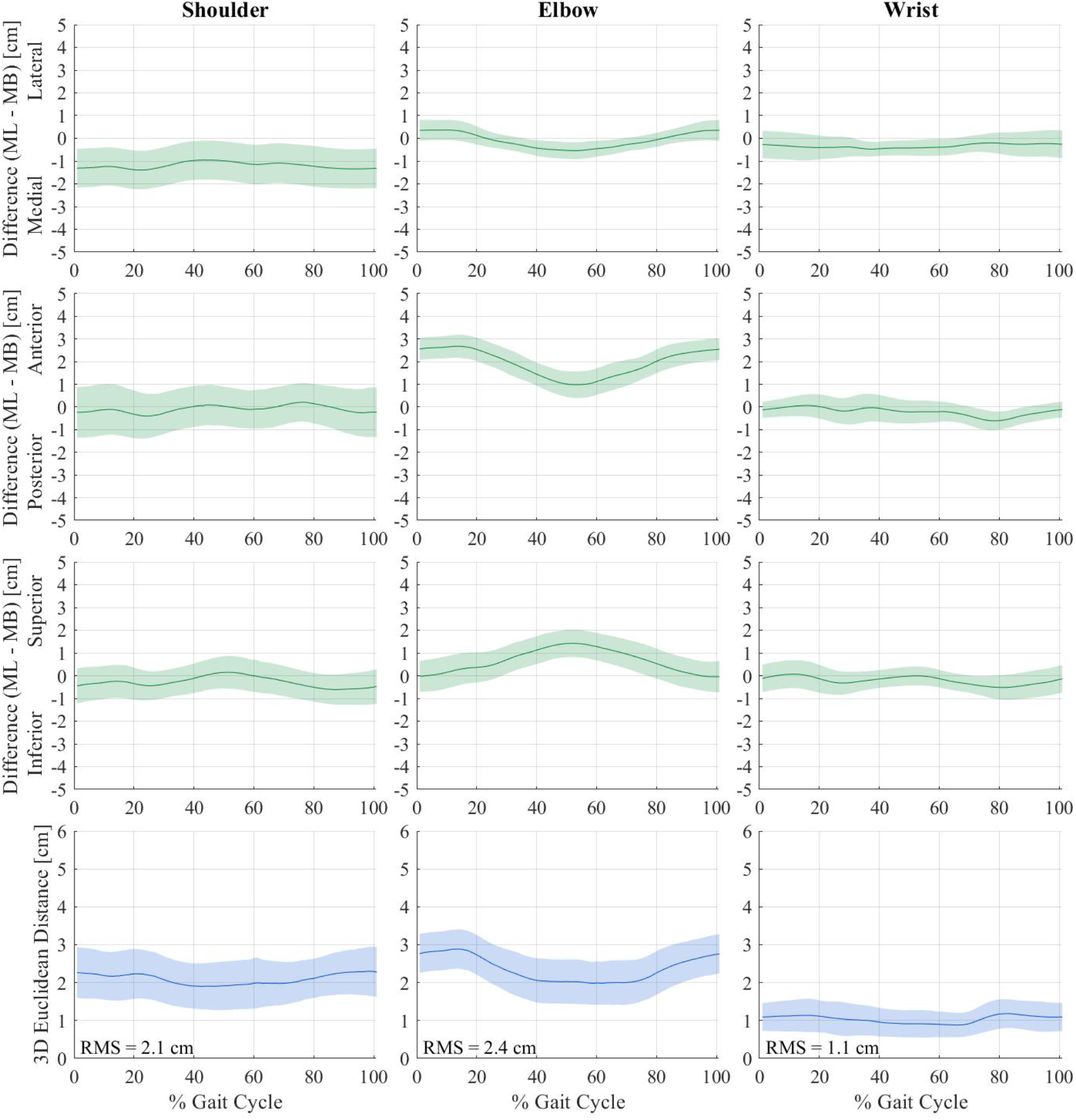
Mean +/− SD upper limb joint position differences between both motion capture systems across the gait cycle for thirty subjects. Differences were calculated as [markerless (ML) position] - [marker-based (MB) position] and are expressed as components along each of the global coordinate system axes (row 1: medial/lateral direction; row 2: anterior/posterior direction; row 3: superior/inferior direction) and as 3D Euclidean distances (row 4). Axis labels (e.g. Lateral) indicate the position of the markerless joint relative to the marker-based joint. Root-mean-square (RMS) values are inset in the 3D Euclidean distance plots.

### 3.2 Lower Limb Global Segment Angles

Subject-average lower limb global segment angles measured using both motion capture systems and the average difference between the measurements are shown in Figure 4. The thigh and shank segments were found to have very similar global segment angles between the marker-based and markerless system for the x- and y-components, with average RMS differences of 0.9°-2.2°. The thigh and shank segment angle z-components had greater differences between the marker-based and markerless systems, of 8.5° and 13° respectively. The foot segment angle x-components capture the greatest range of motion among the lower limb segments, and very similar patterns were measured by both systems across all subjects, with an average RMS difference of 5.4°. The foot segment angle y-components (similar to inversion/eversion) had greater differences between systems with an average RMS difference of 7.0°, while the z-components (similar to toe in/toe out) had little difference with an average RMS difference of 2.8°.

**Figure 4:**
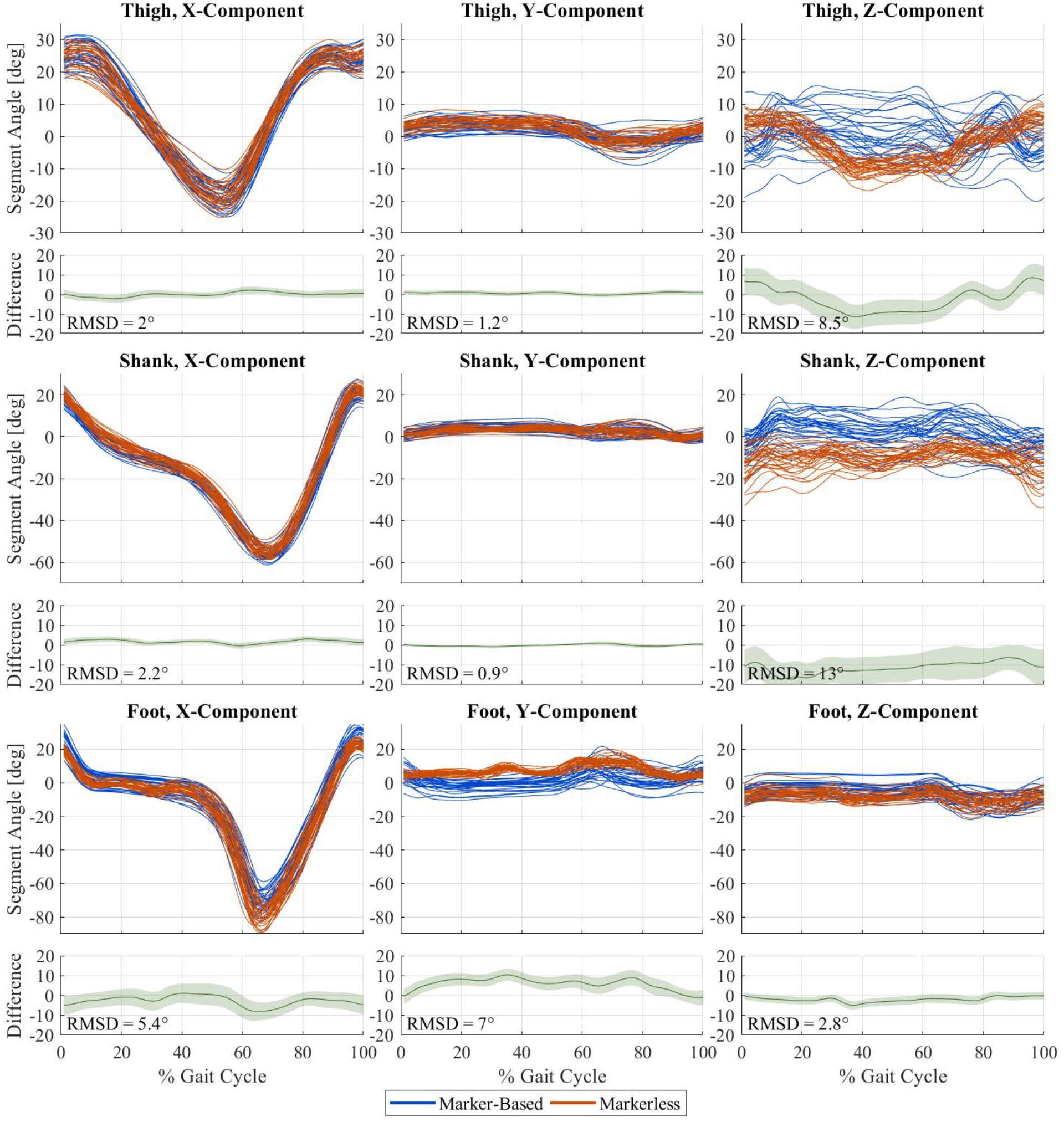
Subject-average global segment angles of the thigh (row 1), shank (row 2), and foot (row 3) for thirty subjects measured by the markerless (orange) and marker-based (blue) motion capture systems, stacked above the average difference (markerless - marker-based) across all subjects. Segment angle x-components represent rotation about the laboratory GCS x-axis, z-components represent rotation about the axially aligned axis of the segment LCS, and y-components represent rotation about the floating axis produced by the cross product of the latter with the former. Average RMS differences are inset in each segment angle difference plot.

### 3.3 Lower Limb Joint Angles

Subject-average lower limb joint angles measured using both motion capture systems are shown in Figure 5, and lower limb joint angles for all cycles from one representative subject are shown in Figure 6. Hip flexion/extension angles showed an offset between systems that decreased during early stance and at toe-off, which resulted in an average RMS difference of 11°. The ankle flexion/extension angles showed a similar but smaller offset that resulted in an average RMS difference of 6.7°. The knee flexion/extension angles demonstrated the greatest similarity among sagittal plane angles, with an average RMS difference of 3.3°. For all three joints, the marker-based system measured greater flexion than the markerless system.

**Figure 5:**
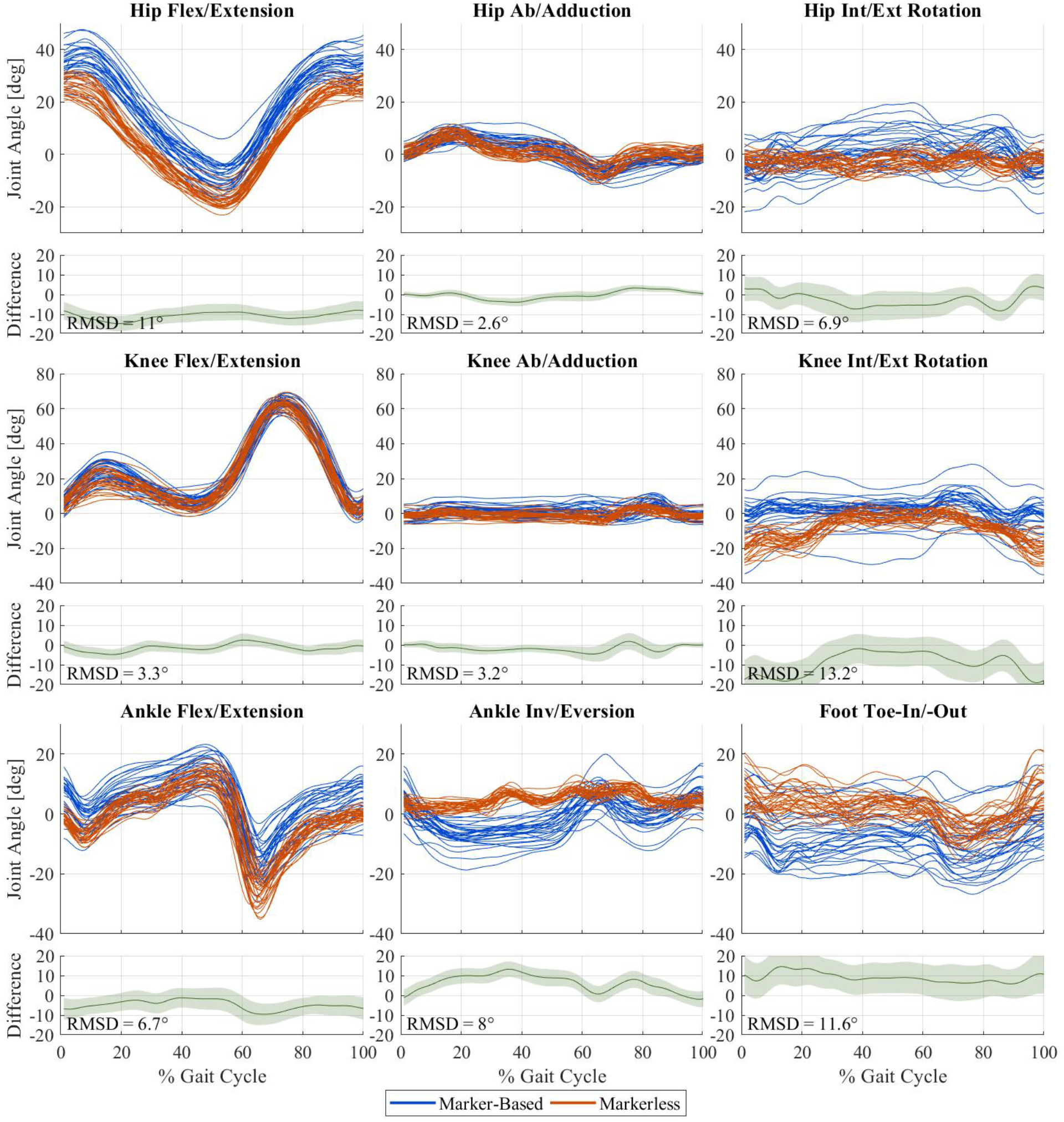
Subject-average joint angles for the hip (row 1), knee (row 2), and ankle (row 3) for thirty subjects measured by the markerless (orange) and marker-based (blue) motion capture systems, stacked above the average difference (markerless - marker-based) across all subjects. Average RMS differences are inset in each joint angle difference plot.

**Figure 6:**
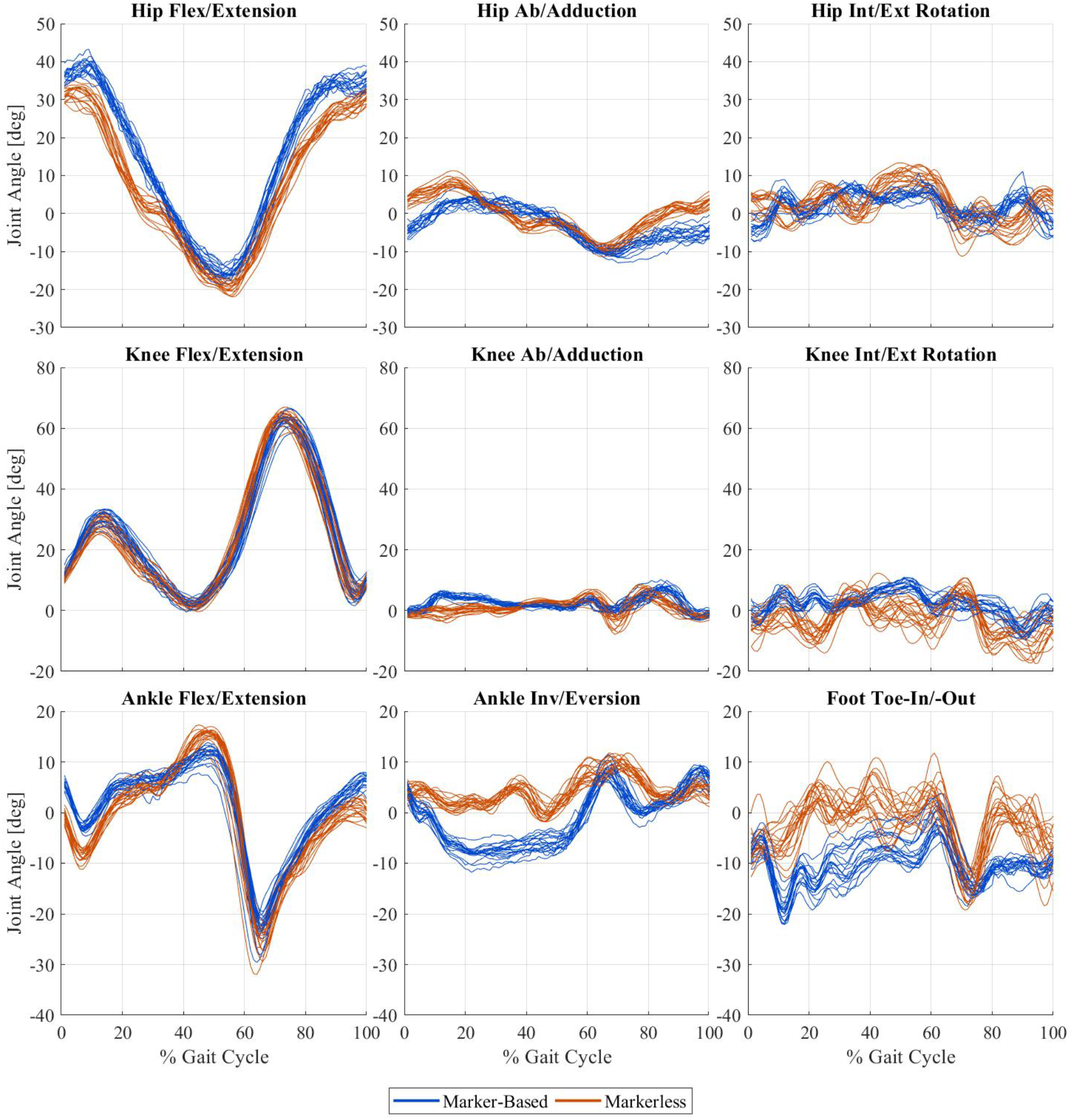
Right lower limb joint angles for 19 gait cycles from one representative subject for the hip (row 1), knee (row 2), and ankle (row 3) measured by the markerless (orange) and marker-based (blue) motion capture systems.

The hip and knee ab/adduction joint angles showed high agreement between the marker-based and markerless systems, which had average RMS differences of 2.6° and 3.2°, respectively. The marker-based system typically measured greater knee adduction than the markerless system, particularly during swing phase (60-100% gait cycle). The ankle inversion/eversion angles had greater differences between systems, particularly during stance phase (10-50% gait cycle); the average RMS difference between systems for this angle was 8.0°.

The hip internal/external, knee internal/external, and foot toe-in/toe-out joint angles measured by the marker-based system had greater variability across all thirty subjects, whereas the joint angle waveforms from the markerless system were more similar across subjects. The internal/external hip rotation angles measured by the markerless system were more neutral than those from the marker-based system, and the average RMS difference between systems was 6.9°. The internal/external knee joint angles differed more during early stance and late swing phases, and the average RMS difference was 13.2°. The foot toe-in/toe-out angles measured by the markerless system follow a similar pattern to those measured by the marker-based system, but with less variability across subjects. The average RMS difference between the foot toe-in/toe-out angles was 11.6°.

## 4. Discussion

This study compared the kinematics of healthy human treadmill gait measured by a marker-based motion capture system and a markerless motion capture system, finding comparable results between the two systems. We have previously shown that spatiotemporal gait parameters are comparable between marker-based and markerless motion capture, using the same dataset analyzed here, and comparable to those from a pressure-sensitive gait mat (Kanko et al., 2021b).

Across all thirty subjects, the joint position estimates differed by less than 3 cm for all joints except the hip, which differed by 3.6 cm. These differences are similar in magnitude to the errors associated with different marker-based joint center location techniques which have been reported up to 4.59 cm for functional methods and 5.08 cm for predictive regression methods (Kainz et al., 2015). The lower limb joint position differences were more sensitive to gait cycle phase compared to the upper limb joints, which may be a result of greater soft tissue artefacts for the lower limbs, less accurate markerless tracking due to occlusion by the treadmill safety bar, or blurred images due to the higher speeds at which the lower limbs travel during walking.

Both systems measured segment poses with similar x-components, which capture the largest segment rotations during gait, differing by roughly 2° for the shank and thigh, and 5.4° for the foot. Shank and thigh segment angle y-components and foot segment angle z-components had similarly small differences of less than 3° between systems. The largest segment angle differences were found for the foot segment angle y-component and the shank and thigh segment angle z-components, which are angular measurements taken about the axially aligned LCS axis during gait. In combination, the relatively small rotations of these segments about these axes and the alignment of the anatomical reference frames between the segments make these movements challenging for both systems to measure, resulting in greater differences of 7°-13°.

The lower limb joint angles captured similar patterns and ranges of motion in flexion/extension at the ankle, knee, and hip, and in ab/adduction at the knee and hip. Joint angles are measured between two adjacent body segments, and therefore any errors in segment pose are amplified in these measurements. The offset in the ankle flex/extension angles is likely a result of the foot segment angle x-component differences, which had an average RMSD of 5.4°. Similarly, the offset in the hip flex/extension angles is likely a result of an offset in the pelvis segment pose between systems, since the differences in thigh segment angle x-components were small, with an average RMSD of 2.0°. Hip and knee internal/external rotation, foot toe-in/toe-out, and ankle inversion/eversion had the most apparent differences between the marker-based and markerless systems, with respect to both the waveform patterns captured and the variability among subjects. The larger differences observed in these joint angles are both notable and challenging to interpret due to the likely presence of errors in the signals from both systems. Therefore, the differences between systems cannot be interpreted as measurement errors of either system, given that the ground truth is not known and both systems likely contribute to the observed differences. We chose to present subject average waveforms and individual gait cycle waveforms for one representative subject to illustrate the differences between systems, capturing both the intrinsic differences in measurement modalities and potential measurement inaccuracies of both systems.

There are several potential sources of error that could affect the measurements from both systems. Marker-based kinematics are susceptible to marker placement variation, kinematic crosstalk, soft tissue artefact, and joint position regression errors. Inconsistent marker placement has been shown to contribute up to 5° of error in lower limb joint angles due to the procedure being performed by different operators and following different protocols (Gorton et al., 2009; Schwartz et al., 2004). To minimize this source of error, markers were placed by the same examiner across all subjects. It has also been shown that skin-mounted marker clusters move relative to the underlying bone during gait, with translations of up to 1.5 cm at the shank and 2.5 cm at the thigh, and rotations up to 8° (Benoit et al., 2015). These tissue artefacts introduce unpredictable, subject- and task-specific errors of up to 3° in knee joint angles (Benoit et al., 2015). Finally, joint position estimation errors have been found to range from 0 cm to 4.6 cm for functional methods and from 0.7 cm to 5.1 cm for predictive regression methods, which can also contribute up to 3° of error in lower limb joint kinematics (Kainz et al., 2015; Leboeuf et al., 2019).

Markerless kinematics may also be affected by several factors. The treadmill on which subjects walked had a safety bar across the front that partially occluded the lower limbs in several camera views (Figure 1). Subjects wore minimal, skin-tight clothing and motion capture markers which were unfamiliar to the neural networks since the training dataset did not include any images of motion capture subjects or markers. Additionally, the video cameras used were necessary for spatial and temporal synchronization with the marker-based system but were not ideal for the markerless system. The video data consisted of 3 Megapixel images recorded at 85 Hz under indoor lighting conditions, producing images that were somewhat low resolution and blurry at times. More appropriate video cameras would provide higher quality images and may improve tracking. Finally, treadmill gait is known to differ from over-ground gait; however, this test compared pose estimation, not characteristics of walking, so any differences are not relevant in the context of this work.

The specific training of the neural networks used for automated pose estimation is both a strength and limitation of the markerless system. Any omissions or biases implicit within training sets will be propagated to error when applied to situations where the training was weak. The sensitivity of the markerless motion capture system to various factors including subject characteristics such as age, sex, ethnicity, health status, and clothing have not yet been fully tested, nor has the sensitivity to environmental factors such as lighting, the use of a treadmill, or being in a laboratory environment. These potential sensitivities will require further testing outside of this study since the results may differ for samples with different characteristics. Finally, the markerless system employs a frame-by-frame approach to track subject movement which may introduce greater noise to kinematic measurements of individual gait cycles compared to marker-based motion capture; however, previous work has indicated that this noise may be overcome by the use of multiple gait cycles and that gait kinematics from markerless motion capture are slightly more consistent across multiple sessions compared to marker-based kinematics (Kanko et al., 2021a).

The results presented here indicate that the overall pose of subjects is measured very similarly between systems, with small joint position differences and similar waveform patterns for most joint and segment angle measures. Given the practical benefits of markerless motion capture such as minimal subject preparation time and reduced collection environment restrictions, this technology could allow studies to be undertaken with truly large sample sizes that were previously not feasible. Furthermore, previous work has indicated that this system offers slightly greater repeatability across multiple collection sessions than marker-based systems, increasing the opportunity for and potential of multi-center trials.

## Conflict of Interest Statement

Scott Selbie is the CEO of Theia Markerless Inc. (Kingston, Ontario), the developers of *Theia3D*.

## Acknowledgements

At the time of the study, RMK was supported by an NSERC Canadian Graduate Scholarship. We thank Human Mobility Research Laboratory members with participant recruitment, data collection, and data processing.

## Notes

### Competing Interest Statement

WSS is the CEO of Theia Markerless Inc. (Kingston, Ontario), the developers of Theia3D. He contributed to the conception and design of the study, critically revised the article for intellectual content, and provided final approval of the submitted version. WSS was not involved with the collection, analysis, or interpretation of data.

